# Delayed start of first-time breeding and non-breeders surplus in the Western Siberian population of the European Pied Flycatcher

**DOI:** 10.1101/387829

**Authors:** Vladimir G. Grinkov, Helmut Sternberg

## Abstract

Non-breeders are those sexually mature individuals that do not breed in a given reproductive cycle of a population. There is a widespread belief that the presence of non-breeders can affect the actual population dynamics, as well as the population responses to environmental change (Lee et al. 2017). Sternberg (1989), using demographic data, has shown that 83% and 62% of males and females, respectively, do not breed in the first year of life in the Lower Saxony (Germany) population of the European Pied Flycatcher (*Ficedula hypoleuca*). Later, with experimental removal of males, it has been proven that in the Lower Saxony and Moscow Region (Russia) populations, there are many non-breeding males (Sternberg et al. 2002). For the Netherlands population of the European Pied Flycatcher, the presence of a large number of non-breeders has been demonstrated using experimental removals for both males and females (Both et al. 2017). Here we have estimated the number of non-breeders in the Western Siberian population of the European Pied Flycatcher using demographic data (11 cohorts from 2001 to 2011 of birth) and experimental removal of males. We have shown that both males and females can start to breed at the age of one to five years. The proportion of non-breeders can be 59.5% and 68.5% for first-year males and females, respectively. We discuss the differences in the proportion of non-breeders between the Western Siberian and European populations of the European Pied Flycatcher, as well as factors affecting the number of non-breeders.

## INTRODUCTION

### Definition

Non-breeders are those sexually mature individuals that do not breed in a given reproductive cycle of a population.

### Importance

Population models that do not explicitly incorporate non-breeders give upwardly biased estimates of the deterministic population growth rate and demographic variance particularly when the equilibrium ratio of nonbreeders to breeders is high (Lee et al. 2017). Estimates of a population growth rate from empirical observations of breeders are only substantially inflated when individuals frequently re-enter the breeding population after periods of nonbreeding (Lee et al. 2017).

Changes in demographic rates of breeders vs. non-breeders differentially affect the population growth rate. In particular, the population growth rate is most sensitive to non-breeder parameters in long-living species, when the ratio of non-breeders to breeders is positive, and when individuals are unlikely to breed at several consecutive time steps (Lee et al. 2017).

### Life histories of non-breeders

There are several hypotheses suggesting strong differences in proportion and life histories of non-breeders:

1. Many non-breeding birds were believed to have resulted from competition for nest sites (Sternberg 1989). A high number of European Pied Flycatcher non-breeders is actually present in the breeding area, and these birds will be breeding within the same area in future (Sternberg et al. 2002; Both et al. 2017).
2. It may be that some young birds do not leave their wintering quarters (Lundberg & Alatalo 1992).
3. “Non-breeders” breed for the first time in other areas due to dismigration (dispersal/spacing) (Berndt & Sternberg 1968; Sokolov 1999; Sternberg et al. 2002).

### Non-breeders in the European Pied Flycatcher

Using demographic data, it had been shown that 83% and 62% of males and females, respectively, do not breed in the first year of life in the Lower Saxony (Germany) population of the European Pied Flycatcher (Sternberg 1989).

Later, with experimental removal of males, it had been proven that in the Lower Saxony and Moscow Region (Russia) populations are indeed many non-breeding males (Sternberg et al. 2002).

For the Netherlands population of the European Pied Flycatcher, the presence of a large number of non-breeders has been demonstrated using experimental removals for both males and females (Both et al. 2017).

### Behaviour of non-breeders in nesting areas

Non-breeders in the European Pied Flycatcher can take part in feeding the nestlings of other pairs of their species (Ilyina 2012). They can participate in extra-pair copulations (Tomotani et al. 2017).

Thus, failing to account for non-breeders in population studies can obscure low population growth rates that should cause management concern. Quantifying the size and demography of the non-breeding section of populations and modelling appropriate demographic structuring is therefore essential to evaluate non-breeders’ influence on deterministic and stochastic population dynamics (Lee et al. 2017).

Even though the European Pied Flycatcher is the one of the most studied species of birds, some aspects of its biology are still poorly understood. For example, the behaviour of non-breeding birds has not been studied.

Here we have estimated the number of non-breeders in the Western Siberian population of the European Pied Flycatcher using demographic data and experimental removal of males. We also compared the estimates of non-breeders’ proportions for the two populations studied (the Lower Saxony (Germany) and the Western Siberian (Russia)).

## METHODS

### Species

We conducted research on the European Pied Flycatcher (Ficedula hypoleuca) (Photo 1). This is a small passerine bird in the Old-World flycatcher family. This is a common and widespread species of hollow nesting birds. The species breeds in the forests of Europe, and in the east of the breeding area inhabits the small-leaved forests of Western Siberia. The species winters in subequatorial Africa.

**Photo 1.**
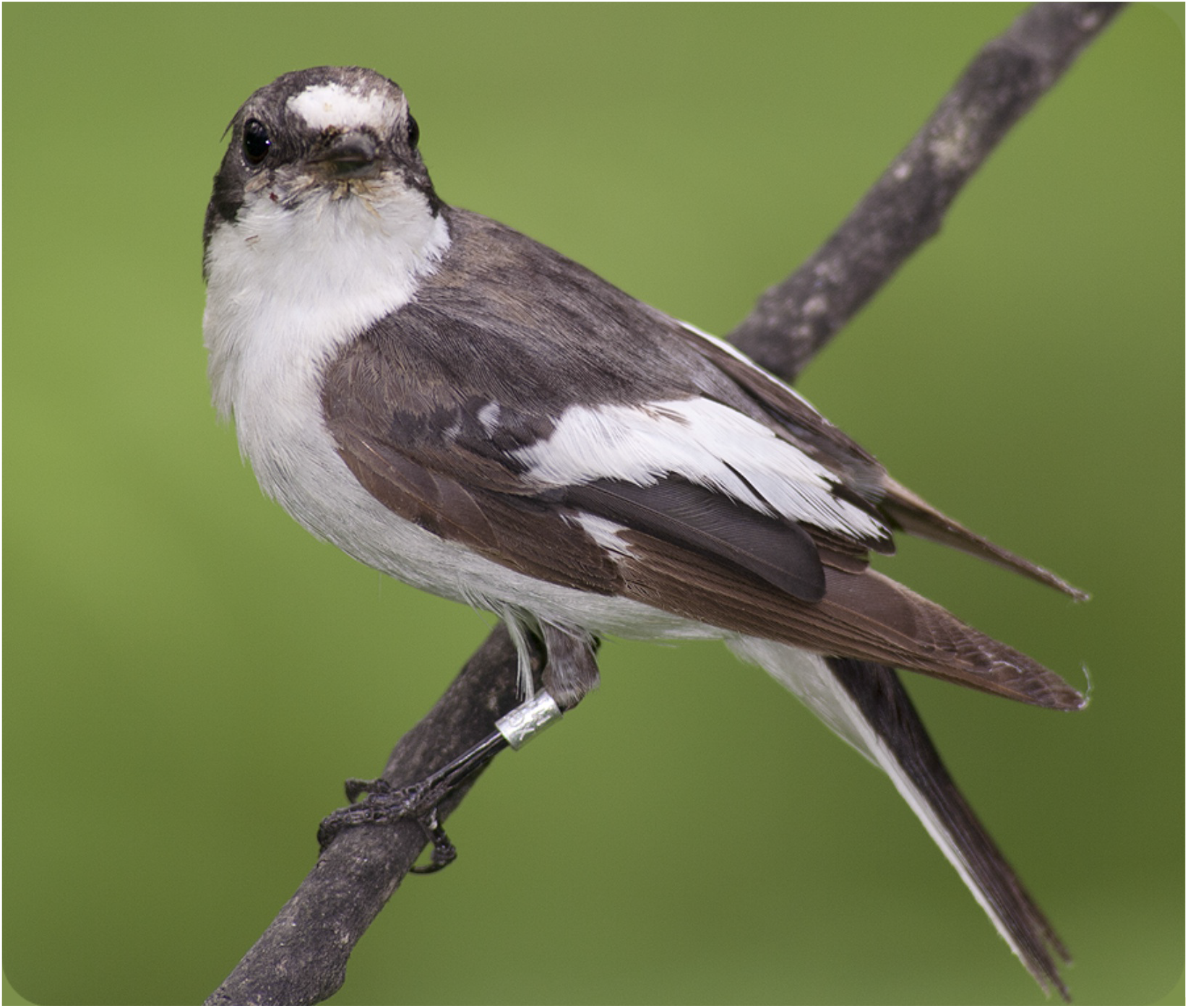
The male of the European Pied Flycatcher

### Place

The study was conducted at the eastern boundary of the European Pied Flycatcher range in Western Siberia in Russia. The nest-box areas were located 13 km south of Tomsk (56°21’N 84°56’E) in a mixed forest.

### Demographic data

The results presented in this poster were obtained during a long-term study of a European Pied Flycatcher population in Western Siberia in the Tomsk region (Bushuev et al. 2012; Kerimov et al. 2014). The study area consisted of three subplots – two 5 ha and one 20 ha areas equipped with nest-boxes. The distance between the most separated nest-boxes in the studied area was approximately 3 km. We have estimated the number of non-breeders in the Western Siberian population of the European Pied Flycatcher using demographic data for 11 cohorts born between 2001 and 2011. This analysis is based on 1089 breeding birds ringed as nestlings whose ages were known (568 males and 521 females).

### Removal experiments

The experiments were conducted in 2002 and in 2004. We divided the study plot into two parts – experimental and control consisting 100 nest-boxes each. In an experimental plot we daily captured all males on arrival and removed them into the keeping aviary. We fed males with mealworms, which were – as well as water – supplied ad libitum. The design of the experiment and aviaries equipment were the same as it has been described in our previous publication (Sternberg et al. 2002). All males were kept in the aviaries until the food in stock was exhausted, and after that all captured males were released at the same time. Later in each breeding season, we tried to recapture experimental males in nests while they were feeding the nestlings to discover which males had successfully paired with females.

### Statistical analysis and software

To calculate the required population parameters, we used the Cormack-Jolly-Seber (CJS) model implemented in the MARK program (White & Burnham 1999) version 9. We used the Akaike Information Criterion (AIC) to select the most parsimonious model and to reveal the most fitting model. For other statistical analyses, we used the R-statistics program (Version 3.5.0 - Copyright © 2018 The R Foundation for Statistical Computing) under an integrated development environment for R - RStudio (RStudio Desktop Version 1.1.447 – Copyright © 2009-2018 RStudio, Inc). To create the poster, we used the program Inkscape™: Open Source Scalable Vector Graphics Editor (Version 0.92.3 – Copyright © 2018 The Inkscape Project, https://inkscape.org) and GNU Image Manipulation Program (GIMP) (Version 2.10.4 - Copyright © 1995-2018, https://www.gimp.org/). To create the distribution version of the poster we used Scribus: Open Source Desktop Publishing (Version 1.4.7 - Copyright © 2015-2016 The Scribus Team, https://www.scribus.net/).

## RESULTS

### Demographic data

#### Females

Females can start breeding at the age of one to five years (Fig. 1). Only 33.0% of all returned females begin to breed from the first year of life. Consequently, 67% of the returned females are non-breeding or pre-breeders in the first year of life. In Lower Saxony 62% of yearling females did not breed. The estimated recruitment rate of females is 12.8% for the Western Siberian population.

**Figure 1.**
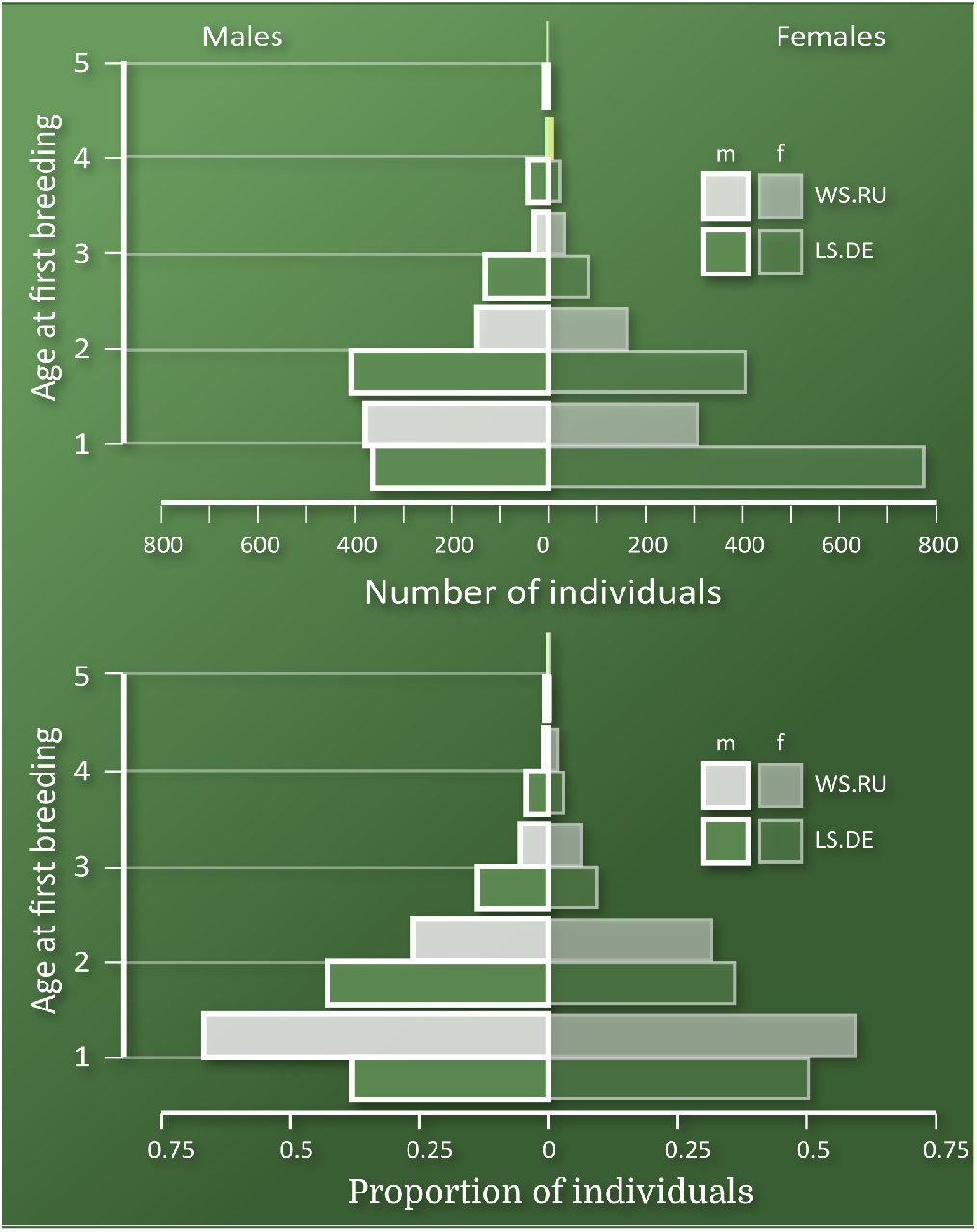
Age at first breeding in Lower Saxony (LS.DE) and Western Siberia (WS.RU)

Using the MARK and Cormack-Jolly-Seber (CJS) model, we calculated that about 62% of females do not breed in the Western Siberian population and 57% in Lower Saxony in their first year of life (Fig. 2, the encounter probability p for transition Pull −> 1 corresponded with the proportion of breeding birds in their first year of life, and 1-p equals to the proportion of non-breeders). The females’ apparent survival between birth and the first year of life (the recruitment rate) in the Western Siberian population was 11.1% (according to the CJS model estimates), in Lower Saxony this figure was 13.4% (Fig. 2, apparent survival Phi for transition Pull −> 1 corresponded with the return rate).

**Figure 2.**
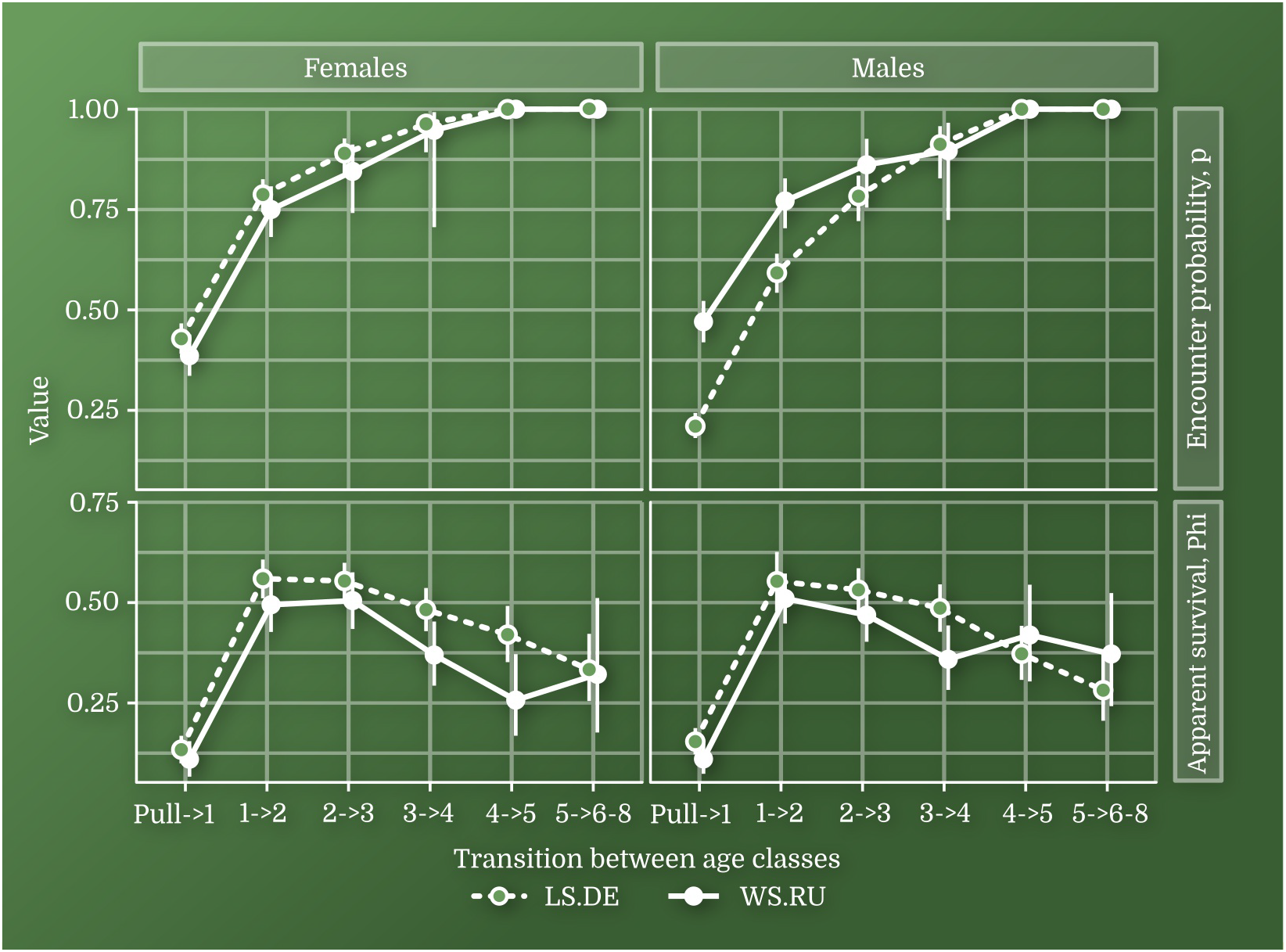
Encounter probability (p) and apparent survival (Phi) calculated for locally born birds in Lower Saxony (LS.DE) and Western Siberia (WS.RU). Vertical bars showed 95% confidence interval

#### Males

Males can start breeding at the age of one to five years. 42.3% of all returned males begin to breed from the first year of life (Fig. 1). Consequently, 57.7% of the returned males are non-breeders in the first year of life (in Germany 83% of yearlings). The estimated recruitment rate of males is 12.3% for the Western Siberian population (Fig. 1).

The CJS model estimated the proportion of non-breeding males in the first year of life at 53% for Western Siberia and 79% for Lower Saxony. Males’ apparent survival between birth and the first year of life (the recruitment rate) was 11.1% for Western Siberia and 15.4% for Lower Saxony (Fig. 2).

The Akaike Information Criterion (AIC) has shown that models in which all parameters are different fit the data best, i.e. there are significant intrapopulation differences in the apparent survival and the encounter probability between the sexes and interpopulation differences in the parameters analysed.

### Removal experiments

#### Proportion of breeders and non-breeders

In 2002 we caught 121 males and released them on 13th May. In 2004 we were able to catch 296 males which were released on 16th May.

There were 4.2 times more males caught during the removal experiment in 2004 in the Western Siberian population, than they were breeding in the control plot. The proportion of non-breeding males in the Western Siberian population may be 69 till 76% (the minimum estimate of the proportion of non-breeders is calculated by the number of nests in the control plot, the maximum by the number of males breeding in the control plot). In Lower Saxony, the proportion was 70% (Sternberg et al., 2002).

In the Western Siberian population, 48.2% of the males caught in the removal experiments started breeding in the year of the removal experiment, which is almost twice as much as in Lower Saxony (X-squared = 31.4, df = 1, p <0.001) (Fig. 3). However, in the following years in the Western Siberian population, only 65% of the caught males started with breeding, and in Lower Saxony the proportion of such birds was 87% (X-squared = 47.9, df = 1, p <0.001) (Fig.3).

**Figure 3.**
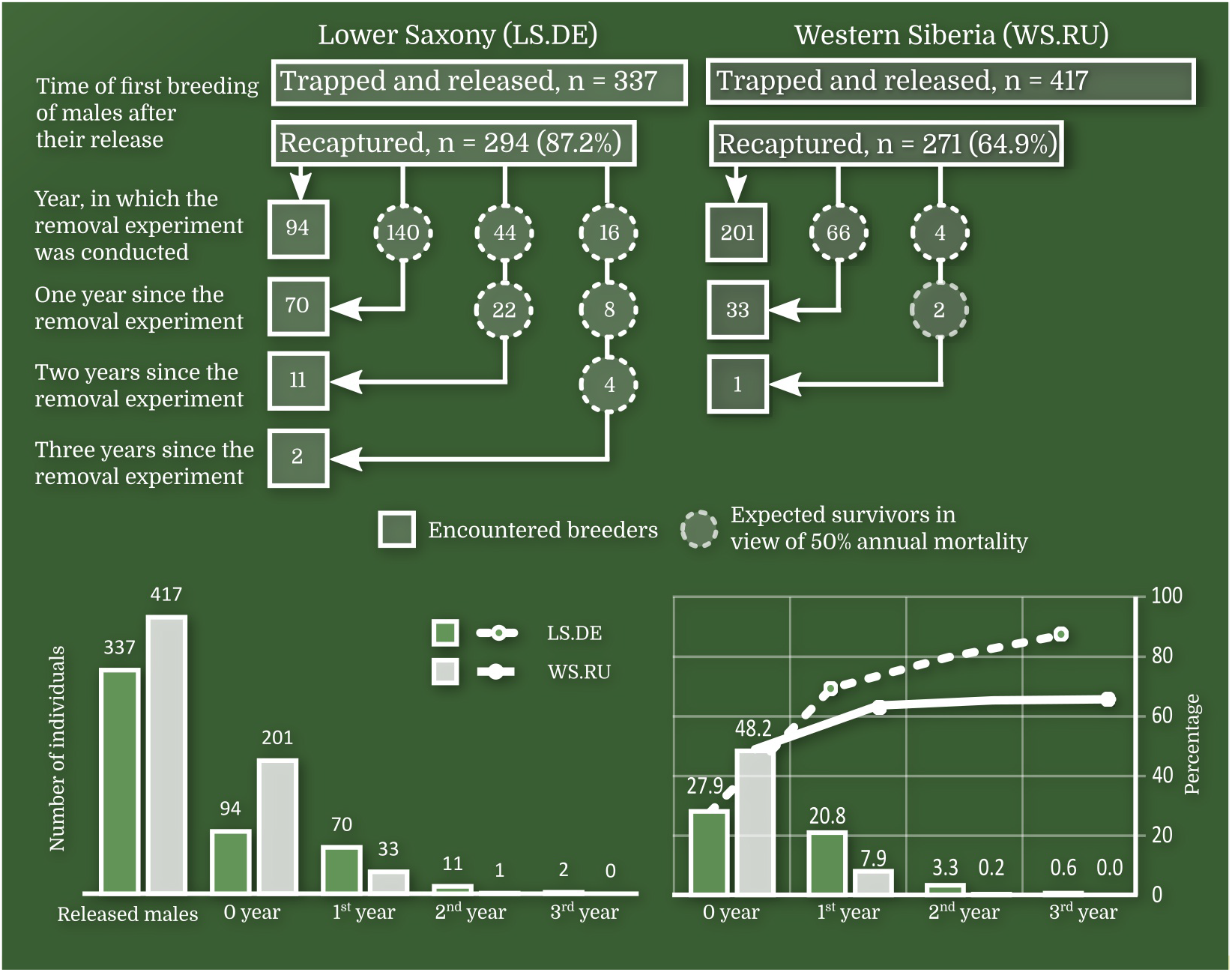
The main results of the removal experiments in Lower Saxony (LS.DE) (taken from Sternberg et al. 2002) and Western Siberia (WS.RU)

#### Features of breeders and non-breeders

We found that older males arrived earlier in Western Siberia in 2004 (Fig. 4) (Kolmogorov-Smirnov Test, p < 0.001 for age groups 1 and >=2; Kolmogorov-Smirnov Test, p < 0.001 for age groups “x” and “x+1”) and they more often acquired a mate (rs = 0.26, n = 417, p < 0.001). The males which had nests in previous breeding seasons (experienced males) more often successfully attracted a female (rs = 0.27, n = 417, p < 0.001). Because breeding experience and age are functionally related, i.e. there are no experienced yearlings in the population, we restricted the sample to old males. Among the latter experienced males were still more successful in attracting females (rs = 0.23, n = 71, p <0.05). Also, yearlings with heavier weight had more chances to get females (rs = 0.12, n=322 p < 0.05). We could not find this correspondence among old males.

**Figure 4.**
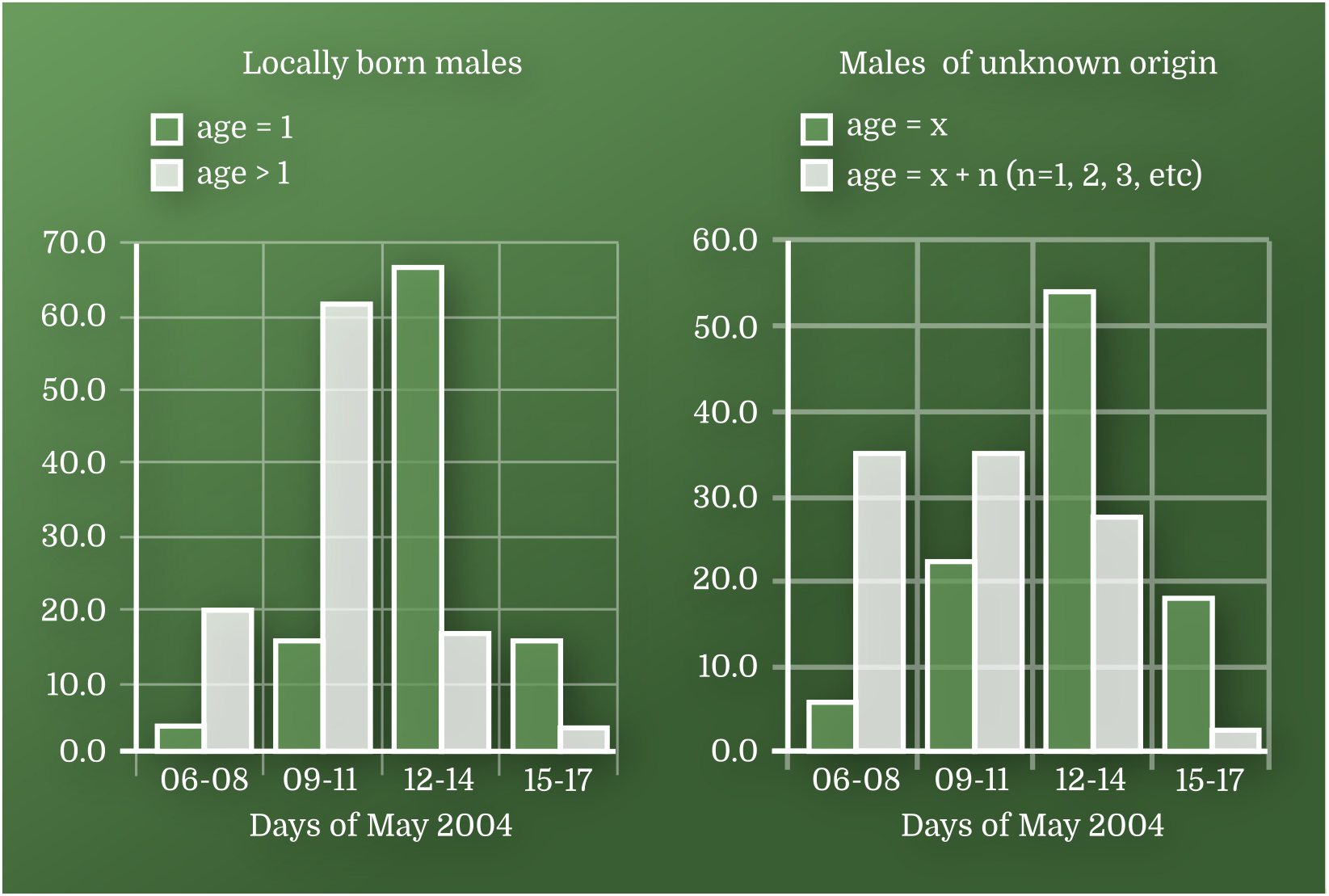
The proportion of different age groups of males caught in different periods of the removal experiment in 2004 (Western Siberia, Russia)

### The other important differences between the populations studied

In both investigated populations, the level of nest predation during the breeding season (for various reasons – from eggs’ and nestlings’ predation to the disappearance of parents) was similar and did not exceed 20%.

The populations under study differ significantly in the mean lifetime number of fledglings (Fig. 5).

**Figure 5.**
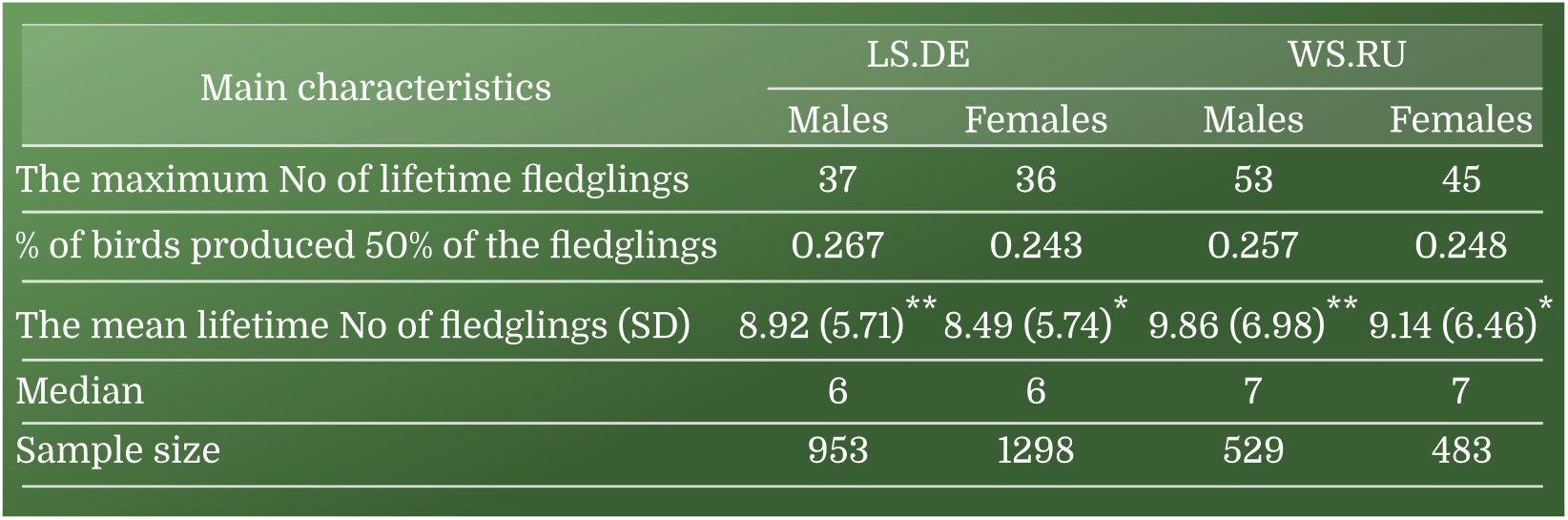
The lifetime reproductive success (LRS) of the European Pied Flycatcher in Lower Saxony (LS.DE) and Western Siberia (WS.RU). * the differences between the means for females from the LS.DE and WS.RU populations are significant at p <0.05; ** the differences between the means for males from the LS.DE and WS.RU populations are significant at p <0.01

Interspecific competition for breeding sites is practically absent in the Western Siberian population of the European Pied Flycatcher. Here, the occupation rate of the nest-boxes by European Pied Flycatchers can reach 96% in some years. In Lower Saxony, the occupation rate of nest-boxes in the study sites with European Pied Flycatchers did not exceed 50%, the other half of the nest-boxes were mostly occupied by different species of tits (Parus spp.). Apparently, the almost complete absence of interspecies competition for breeding sites in the Western Siberian population leads to the fact that both males and females are more easily entered in the reproductive part of the population (Fig. 6). This leads to the fact that the encounter probability of males and females of a young age in the Western Siberian population is higher than in Lower Saxony (Fig. 6, Fig. 2).

**Figure 6.**
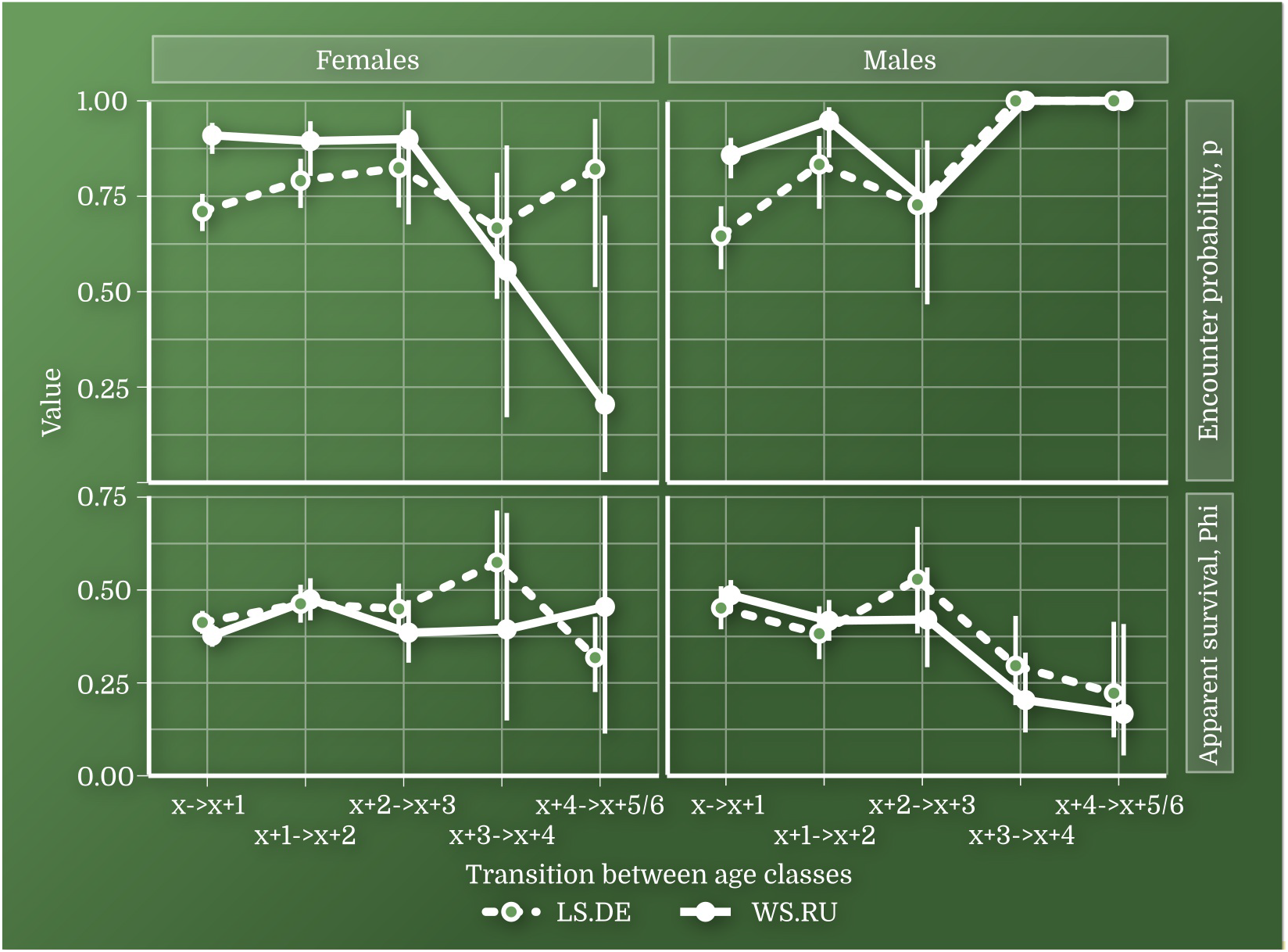
Encounter probability (p) and apparent survival (Phi) calculated for birds of unknown origin in Lower Saxony (LS.DE) and Western Siberia (WS.RU). Vertical bars showed 95% confidence interval

The migration route for birds from the Western Siberian population is almost twice as long as for birds from Lower Saxony (Fig. 7).

**Figure 7.**
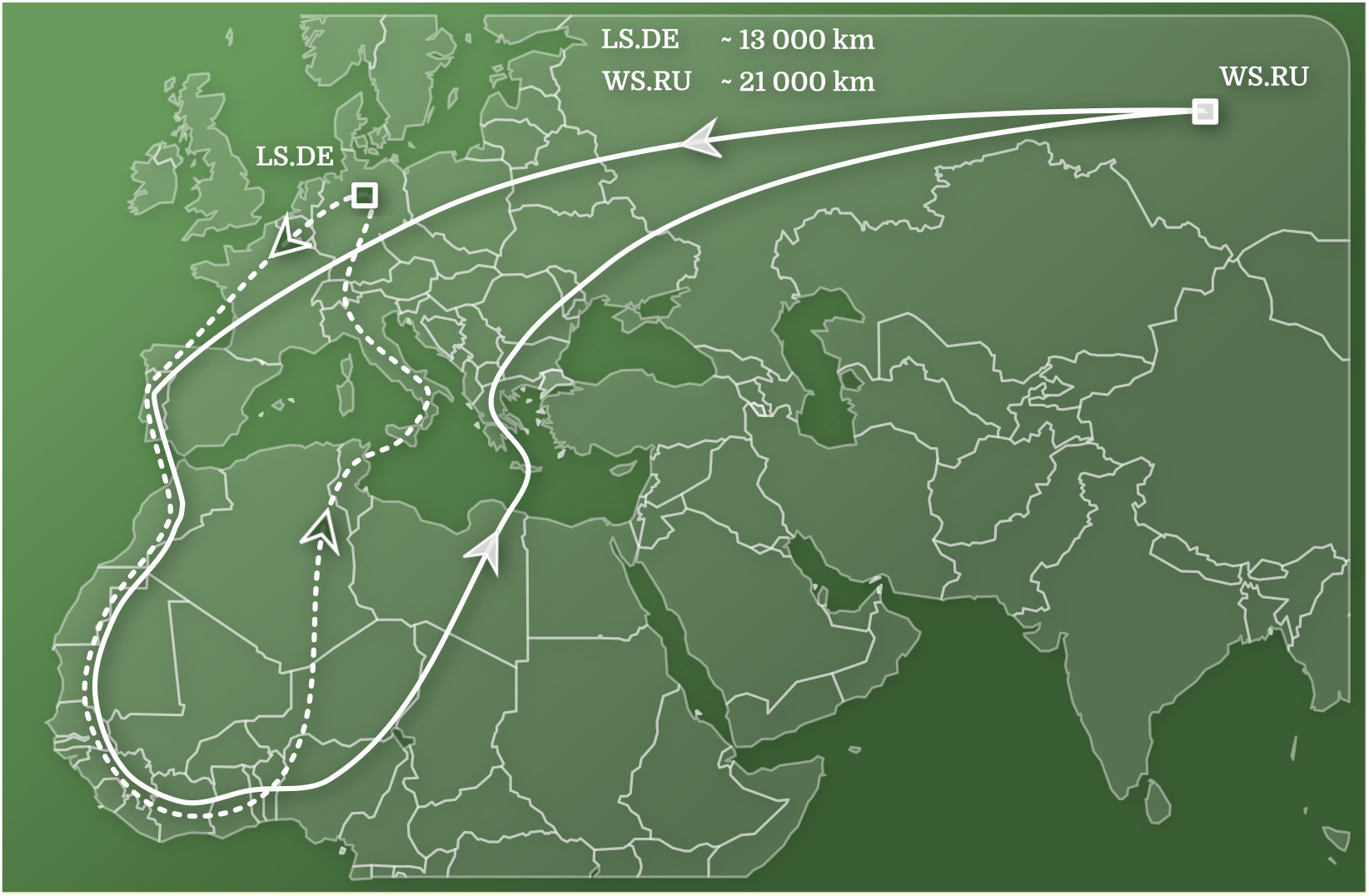
The location of the populations studied and the estimated migration routs of the European Pied Flycatcher for both populations

## CONCLUSION

I. In the Western Siberian population there are differences between males and females in the age of the first breeding: males begin to breed at a younger age. There are also interpopulation differences since males begin to reproduce at a younger age in the Western Siberian population. The females of the Western Siberian population also tend to start reproducing at a younger age than in Lower Saxony. The entry of males from the Western Siberian population into the reproductive part of the population at a younger age may be due to the almost total absence of interspecific competition.
II. The age of the first reproduction correlates to the results of the removal experiment: if in the population the males start to breed at a younger age, then in the year of the removal experiment more males enter the reproductive part of the population.
III. In the two populations studied, there is a population reserve, which is several times larger in number than the reproductive part of the population.
IV. Inter-population differences in the mean lifetime number of fledglings are apparently due to a longer migration of birds from the Western Siberian population. A larger mean lifetime number of fledglings in the Western Siberian population seem to compensate for higher migration costs of this population. It is likely that the absence of interspecies competition for nesting sites in the Western Siberian population allows birds to successfully grow nestlings more often during a lifetime, which leads to a bigger number of grown nestlings for a lifetime.

## ACKNOWLEDGMENTS

This work was supported by Russian Fund of Basic Research RFBR (projects 05-04-49173-a, 06-04-49082-a, 09-04-00162-a, 13-04-01309-a, 18-04-00536-a), and the State Assignment Ch. 2 CITIS AAAA-A16-116021660.

## REFERENCES

Berndt R, and Sternberg H. 1968. Terms, studies and experiments on the problems of bird dispersion. Ibis 110:256–269.

Both C, Burger C, Ouwehand J, Samplonius JM, Ubels R, and Bijlsma RG. 2017. Delayed age at first breeding and experimental removals show large non-breeding surplus in Pied Flycatchers. Ardea 105:43–60.

Bushuev AV, Husby A, Sternberg H, and Grinkov VG. 2012. Quantitative genetics of basal metabolic rate and body mass in free-living pied flycatchers. Journal of Zoology 288:245–251.

Ilyina TA. 2012. Phenomenon of visiting in the pied flycatcher (Ficedula Hypoleuca Pall., Passeriformes, Aves) in the breeding period. Moscow University Biological Sciences Bulletin 67:88–92.

Kerimov AB, Grinkov VG, Ivankina EV, Ilyina TA, and Bushuev AV. 2014. The Influence of Spring Temperature on the Intensity of Advertising Behavior and Basal Metabolic Rate in Bright and Pale Pied Flycatcher (Ficedula Hypoleuca) Males. Zoologichesky Zhurnal 93:1288–1302.

Lee AM, Reid JM, and Beissinger SR. 2017. Modelling effects of nonbreeders on population growth estimates. Journal of Animal Ecology 86:75–87.

Lundberg A, and Alatalo RV. 1992. The Pied Flycatcher. London: T & AD Poyser ltd.

R Core Team (2018). R: A language and environment for statistical computing. R Foundation for Statistical Computing, Vienna, Austria. https://www.R-project.org/.

R Studio Team (2016). RStudio: Integrated Development for R. RStudio, Inc., Boston, MA URL http://www.rstudio.com/.

Sokolov L. 1999. Philopatry, dispersal and population structure of passerines on the Courish Spit of the Baltic Sea. Vogelwarte: Zeitschrift für Vogelkunde 40:302 – 314.

Sternberg H. 1989. Pied flycatcher. In: Newton I, ed. Lifetime reproduction in birds. london: Academic Press Inc, 55–74.

Sternberg H, Grinkov VG, Ivankina EV, Ilyina TA, Kerimov AB, and Schwarz A. 2002. Evaluation of the size and composition of nonbreeding surplus in a Pied Flycatcher Ficedula hypoleuca population: Removal experiments in Germany and Russia. Ardea 90:461–470.

Tomotani BM, Caglar E, de la Hera I, Mateman AC, and Visser ME. 2017. Early arrival is not associated with more extra-pair fertilizations in a long-distance migratory bird. Journal of Avian Biology 48:854–861.

White GC, and Burnham KP. 1999. Program MARK: survival estimation from populations of marked animals. Bird Study 46: S120–S139.

